# Local opposite orientation preferences in V1: fMRI sensitivity to fine-grained pattern information

**DOI:** 10.1101/036269

**Authors:** Arjen Alink, Alexander Walther, Alexandra Krugliak, Nikolaus Kriegeskorte

## Abstract

The orientation of a visual grating can be decoded from human primary visual cortex (V1) using functional magnetic resonance imaging (fMRI) at conventional resolutions (2-3 mm voxel width, 3T scanner). It is unclear to what extent this information originates from different spatial scales of neuronal selectivity, ranging from orientation columns to global areal maps. According to the global-areal-map account, fMRI orientation decoding relies exclusively on fMRI voxels in V1 exhibiting a radial or vertical preference. Here we show, by contrast, that 2-mm isotropic voxels in a small patch of V1 within a quarterfield representation exhibit reliable opposite selectivities. Sets of voxels with opposite selectivities are locally intermingled and each set can support orientation decoding. This indicates that global areal maps cannot fully account for orientation information in fMRI and demonstrates that fMRI also reflects fine-grained patterns of neuronal selectivity.

**Significance statement:** Conventional (3T) *functional magnetic resonance imaging (fMRI) allows one measure brain activity at a spatial resolution of 2-3 mm. Brain response patterns in the primary visual cortex (V1) measured with fMRI have been shown to contain robust information about the orientation of visual grating stimuli. However, it is unclear whether this information arises only from global-areal patterns or also from more fine-grained patterns. Here we show that opposite orientation preferences are present and replicable within small V1 patches. This finding demonstrates that fine-grained fMRI patterns contribute to the orientation information present in fMRI data*.

---

Visual orientation is known to be represented in columnar preference patterns in the primary visual cortex (V1) at a sub-millimetre scale (Yacoub et al., 2008). Kamitani and Tong (2005) demonstrated that fMRI patterns measured in V1 at standard resolution (3-mm isotropic voxels) provide information about the orientation of visual gratings. This study had a big impact in part because it suggested a sensitivity of standard-resolution fMRI to columnar-scale neuronal selectivity patterns. However, it has been proposed that V1 orientation decoding might rely on coarse-scale organisations instead (Op de Beeck et al. 2010). In particular, several studies demonstrated slight preferences for radial orientations (Sasaki et al., 2006; Mannion et al., 2010; Freeman et al. 2011; Alink et al., 2013), which might explain orientation decoding results. A left-tilted diagonal grating, for example, will have approximately radial orientation in the upper left and lower right quadrants, driving the corresponding quarterfield representations of V1 more strongly than the other two quarterfield representations (Sasaki et al., 2006). It has been argued that this effect completely explains fMRI orientation decoding (Freeman et al. 2011).

One way to minimize a contribution to orientation decoding from the radial-preference map is to use logarithmic spiral stimuli. A logarithmic spiral has a constant orientation relative to the radial direction, e.g. 45°. Two spirals with orientations 45° and -45°, respectively, relative to the radius are orthogonal to each other everywhere. They are also balanced about the radial direction everywhere, and thus radial preference cannot account for their decodability. However, such spirals have been shown to be robustly decodable (Mannion et al. 2010; Alink et al., 2013; Freeman et al., 2013). In addition to radial-preference, however, there is evidence that V1 patches also respond preferentially to vertical orientation (Mannion et al., 2010; Alink et al. 2013; Freeman et al., 2013). This global vertical preference predicts distinct global-areal patterns to be elicited by opposite-sense spirals and, thus, spiral decoding as well might be explained by global-areal-scale pattern information.

The aim of the current study is to test if fMRI response patterns with a grain finer than these two coarse-scale preference maps contribute to orientation decoding. The observation that orientation decodability is robust to high-pass filtering of fMRI patterns has been considered as evidence for a fine-grained contribution to fMRI orientation decoding (Swisher et al., 2010; Shmuel et al., 2010; Alink et al., 2013). Filtering analysis, however, is not able to conclusively determine whether finegrained activation patterns contribute to orientation decoding because coarse-scale neural effects can give rise to spurious high-spatial frequency fMRI pattern information if adjacent voxels have different sensitivity to local neural activity. This effect is illustrated in Figure 1. Differences in sensitivity (the voxel gain field) can result, for example, from partial volume sampling, with some voxels sampling mainly gray matter and others mainly white matter. A voxel gain field is not expected to invert the sign of a contrast between two stimuli. Therefore, if orientation decoding of gratings and spirals originated solely from coarse-scale radial and vertical preferences, respectively, then one would not expect voxels in a local cluster to exhibit reliable opposite preferences. Under the global areal account of grating decoding (i.e. radial preference), a small patch of V1 representing a region within one visual quarterfield should not contain voxels preferring tangential over radial stimuli. Similarly, under the global areal account of spiral decoding (i.e. vertical preference), a small patch of V1 representing a region within one visual quarterfield should not contain voxels preferring horizontal over vertical stimuli. Here we show that local voxel clusters in V1 do exhibit reliable preferences for both radial and tangential orientations (in the gratings scenario) and for both vertical and horizontal orientations (in the spirals scenario). The opposite preferences are intermingled within small patches of V1, forming a fine-grained pattern. Gratings can robustly be decoded using either only the radial-preferring or only the tangential-preferring voxels. Similarly, spirals can be decoded using either only the vertical-preferring or only the horizontal-preferring voxels. These results clearly demonstrate the reliable presence of voxels of opposite selectivity within local small patches of V1. Fine-grained fMRI patterns, thus, contribute to orientation decoding.

**Figure 1.**
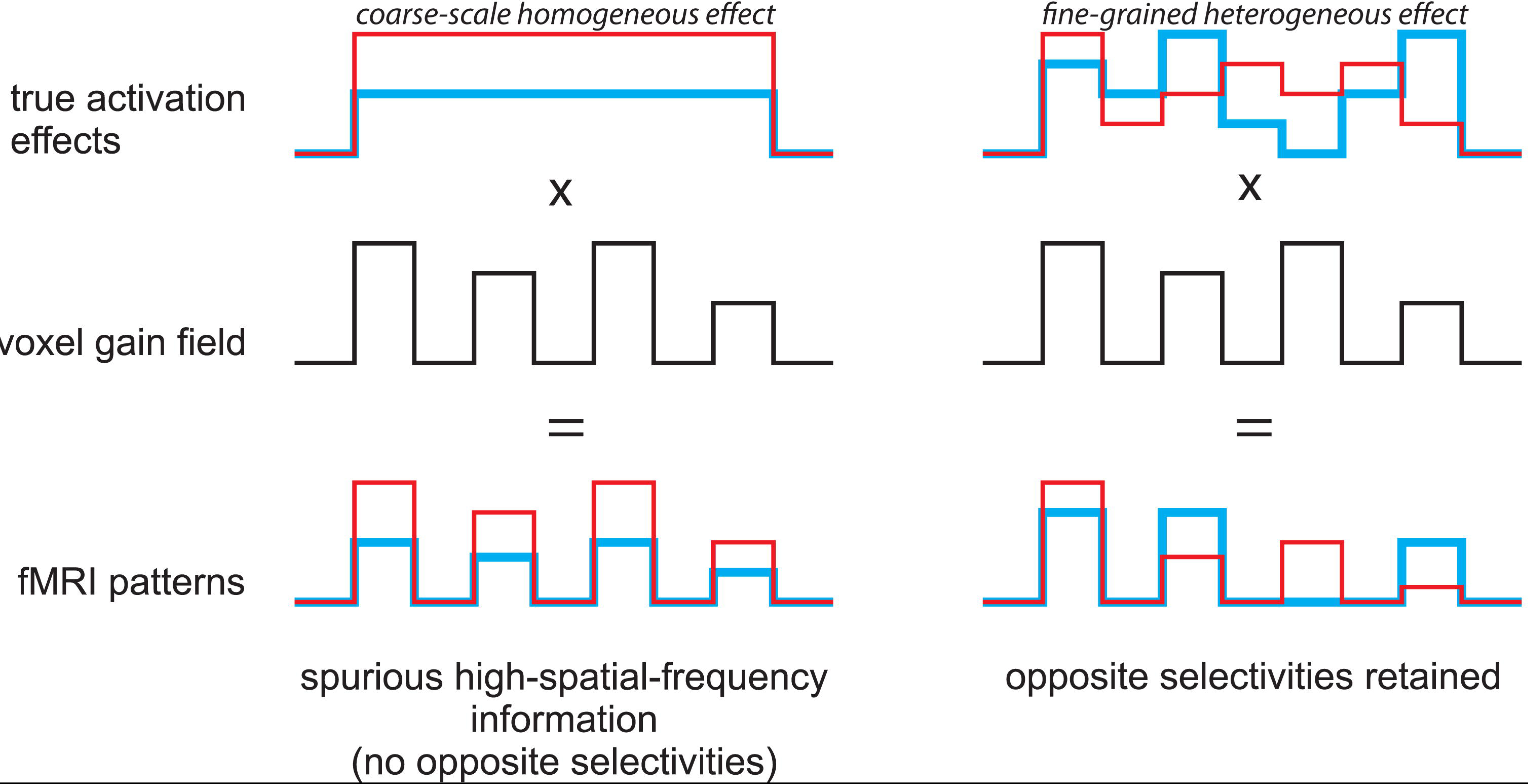
Coarse-scale neural effects can give rise to spurious high-spatial frequency fMRI pattern information in the presence of a high-spatial frequence gain field across voxels. An illustration of how differences in sensitivity to local activation across voxels (the voxel gain field) can lead to spurious high-spatial-frequency information in fMRI patterns. The left column shows the effect of a gain field on a coarse scale homogenous effect and the right column shows the effect of gain field on a fine-grained heterogeneous effect. An important property of the gain field effect is that the signs of the true activation effects are preserved. Spatial filtering analyses will suggest high-spatial frequency information in either scenario (left and right). However, the signature of a finegrained heterogeneous effect (right) is the presence of local opposite selectivities (right only). Note that in actual data the sign of effects can be inverted by fMRI noise; this effect is not illustrated in this figure.

## Materials and methods

### Stimuli and design

*Common features of all stimuli*. All stimulus types were presented within an annulus (inner radius = 1.5°, outer radius = 7.04°) centered on fixation on a mid-gray background. The annulus was divided into 36 log-polar tiles defined by twelve radial lines emanating from the center at 30° offsets and two concentric divisions exponentially spaced between the inner and outer radii (radii including inner and outer: 1.50°, 2.51°, 4.20°, 7.04°). This log-polar tiling was apparent in the form of mid-gray “grout lines” present in all stimuli. For each stimulus type there were two exemplars, which had 90° orientation disparity at every location within the annulus. The oriented edges of all stimuli had 100% contrast. The phases of the oriented edges were randomized across presentations of the same exemplar.

*Gratings*. The orientation of the gratings was 45° clockwise and 45° anti-clockwise from the vertical. The gratings had a spatial frequency of 1.25 cycles per visual degree. This spatial frequency drives V1 strongly (Henriksson et al., 2008) and ensures that even the smallest tiles of the log-polar array contains more than a full spatial cycle.

*Spirals*. We used logarithmic spirals whose edges were at a constant angle of +/−45° relative to the radius emanating from fixation. The spiral stimuli had 22 rectangular contrast cycles along the perimeter. This number of cycles along the perimeter was chosen so as to approximately match the spirals' average spatial frequency across radii to that of the uniform gratings. The two spiral exemplars differed in sense: clockwise or anti-clockwise, lending them 90° orientation disparity at every location. Spiral stimuli are radially balanced because clockwise and anti-clockwise spiral stimuli deviate equally (45°), though in opposite directions, from local radial orientations.

*Experimental design*. Stimuli were presented to each subject in a single fMRI session comprising eight scanner runs, each of which lasted eight minutes. During each run, we presented both exemplars of one stimulus type (e.g. clockwise and anti-clockwise spirals). Subjects were presented with two runs for each stimulus type. Each run was divided into four equal subruns. Each subrun contained six stimulus blocks (three blocks for each exemplar, with exemplars alternating across blocks and the leading exemplar alternating across subruns). Each block lasted 14 s and contained phase-randomized versions of a single exemplar. During a stimulus block, 28 phase-randomized versions of the exemplar were presented at a frequency of 2 Hz. The stimulus duration was 250 ms, followed by an interstimulus interval (ISI) of 250 ms, during which only the fixation dot and a tiny task-related ring around it was visible (see Task, below).

*Retinotopic mapping stimuli*. In order to define regions of interest (ROIs) within V1, we presented dynamic grating stimuli designed to optimally drive early visual cortex. Like the main-experimental stimuli, these stimuli were based on a log-polar array (Figure 2), but without the grout lines and with 20 patches per ring. Each patch contained rectangular gratings with a spatial period of one third of the patch's radial width. Grating orientation and phase was assigned randomly to each patch. Over time, the phase of the gratings increased continuously (1 cycle per second) resulting in continuous motion in each patch (in different directions). In addition, the orientation of the grating increased in steps of n/6, once each second, resulting in motion direction changes within patches over time. We used five such stimuli, driving different parts of the retinotopic representations in V1: (1) a horizontal double-wedge stimulus, spanning a polar-angle range of +/−15° around the horizontal meridian, (2) a vertical double-wedge stimulus of the same kind, (3) a stimulus that covered the region driven by the main-experimental stimulus (1.50°-7.04° eccentricity), (4) a 0.5°-wide ring peripherally surrounding the main-experimental stimulus annulus (7.04°-7.54° eccentricity), and (5) a 0.5°-wide ring inside the annulus (1.00°-1.50° eccentricity). Stimuli were presented in 6-s blocks. This block length was chosen to balance temporal concentration (which increases design efficiency for long blocks due to hemodynamic buildup) and stimulus adaptation (which reduces design efficiency for long blocks due to reduced neuronal responses). The five dynamic stimuli and 6-s fixation periods were all presented 20 times each in a random sequence over a single run lasting 12 min.

**Figure 2.**
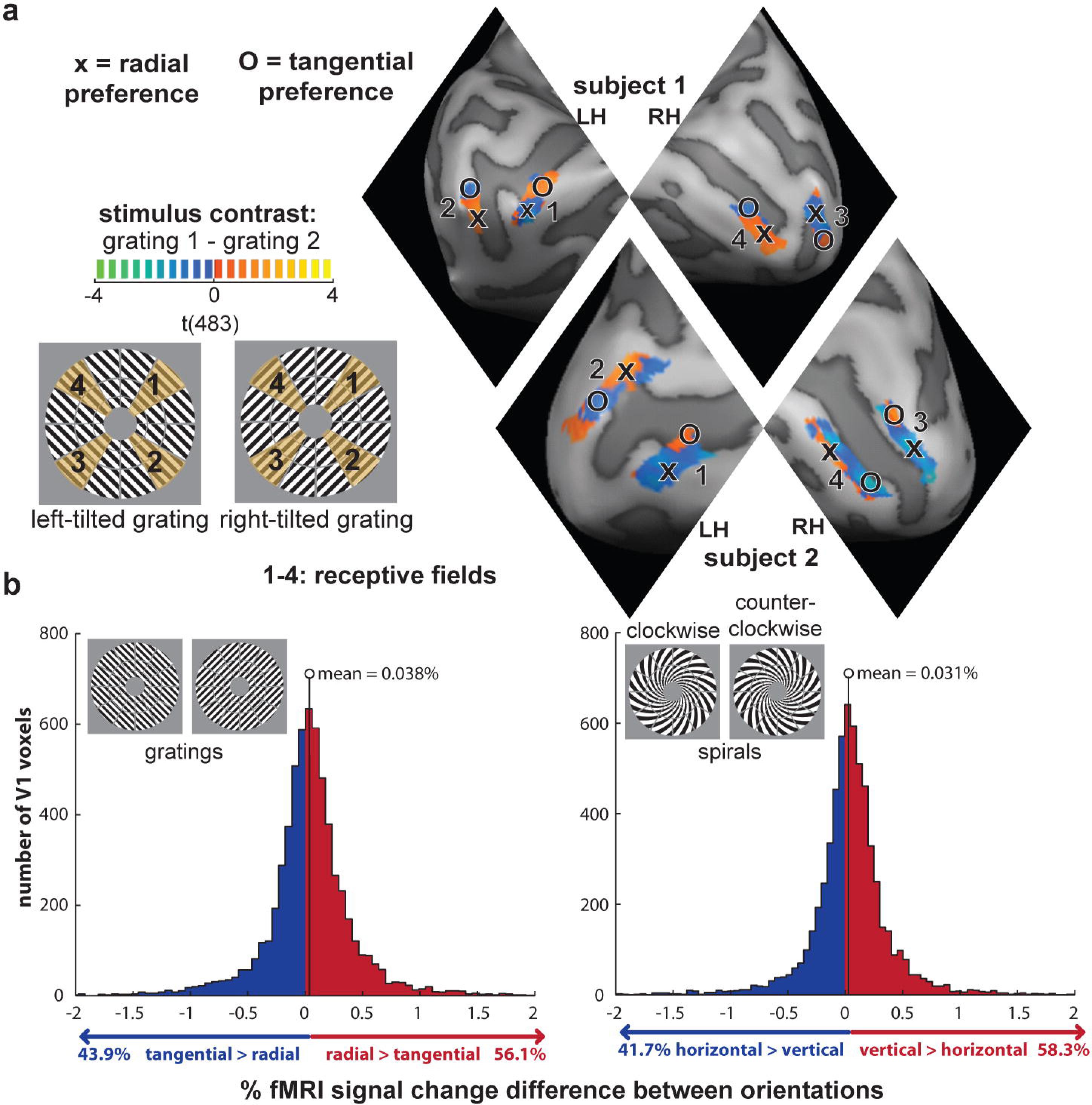
opposite orientation preferences intermingle within quarterfield patches in VI. **(a)** The contrast t maps indicating the activation difference between the two displayed visual gratings. Activation is only shown for the four within-quarterfield ROIs, which are labeled clockwise from 1 to 4. Positive and negative t-values indicate either a radial or a tangential preference depending on the visual field they are in. This we have clarified by labeling local activation clusters with 0 and X when they have a tangential and radial preference respectively. Note that the activation map is based on a conventional GLM t-contrast using all data. **(b)** Histograms showing the distribution of V1 voxels that prefer radial vs tangential orientation (left) and vertical vs horizontal orientation (right). These plots are based on all voxels in all ROIs across all participants.

### Subjects and task

*Subjects*. Eighteen healthy volunteers (13 female, age range 20-39) with normal or corrected-to-normal vision took part in this fMRI experiment. Before the experiment, participants were introduced to the experimental procedure and informed consent was given.

*Task - fMRI*. During all runs, including retinotopic mapping, subjects were instructed to continuously fixate a central dot (diameter: 0.06° visual angle). Centered on the fixation dot, there was a small black ring (diameter: 0.20°, line width: 0.03°), which had a tiny gap (0.03°) either on the left or right side. The gap switched sides at random moments in time at an average rate of once per 3 s (with a minimum inter-switch time of 1 s). The task of the subject was to continuously report the side of the gap by keeping the left button pressed with the right index finger whenever the gap was on the left side, and by keeping the right button pressed with the right middle finger whenever the gap was on the right side. The task served to enforce fixation and to draw attention away from the stimuli.

### MRI measurements and analysis

*MRI measurements*. Functional and anatomical MRI data were acquired with a 3T Siemens Tim-Trio MRI scanner using a 32-channel head coil. During each main run, we acquired 252 volumes containing 31 slices covering the occipital lobe as well as inferior parietal, inferior frontal, and superior temporal regions for each subject using an EPI sequence (TR=2000 ms, TE=30 ms, flip angle=77°, voxel size: 2.0 mm isotropic, field of view: 205 mm; interleaved acquisition, GRAPPA acceleration factor: 2). The same EPI sequence was employed for retinotopic mapping, during which we acquired 360 volumes. For each participant we also obtained a high-resolution (1 mm isotropic) T1-weighted anatomical image using a Siemens MPRAGE sequence.

*Data preprocessing*. Functional and anatomical MRI data were preprocessed using the Brainvoyager QX software package (Brain Innovation, v2.4). The first two EPI images for each run were discarded (affected by T1 saturation effects). After preprocessing (slice-scan-time correction, 3D head-motion correction, linear-trend removal and temporal high-pass filtering removing frequencies below 2 cycles per run), functional data for all subjects were aligned with the individual high-resolution anatomical image and transformed into Talairach space (Talairach & Tournoux, 1988) as a step toward cortex-based analysis in BrainVoyager. After automatic correction for spatial inhomogeneities of the anatomical image, we created an inflated cortex reconstruction for each subject. All ROIs were defined in each individual subject's cortex reconstruction and projected back into voxel space. Note that we did not use Talairach space or a cortex-based common space for ROI definition and within-ROI patterns were analyzed separately in each subject.

*Retinotopic mapping and region of interest definition*. A general linear model (GLM) was fitted to the retinotopic mapping data, with five predictors for the five dynamic grating stimuli based on convolving boxcar functions with the hemodynamic response function as described by Boynton et al. (1996). Activation t-maps for each stimulus type were projected onto polygon-mesh reconstructions of individual subjects' cortices. We determined the borders between V1-2 based on cortical t-maps for responses to vertical and horizontal double-wedge stimuli Regions of interest (ROIs) were only created in the portion of V1 that was more active when presenting the dynamic grating stimulus covering the main-experimental annulus as compared to central and peripheral stimulation. ROIs were defined as patches covering the central third portion of each quarterfield's polar range as visualized in Figure 2. We excluded the remnant of the quarterfield area to reduce spillover of signals between V1 quarterfield representations.

*Pattern-classifier analysis and orientation preference definition*. Preprocessed functional fMRI data for the main experiment and individual ROI coordinates were imported into Matlab using the NeuroElf Toolbox v0.9c (developed by Jochen Weber, Columbia University). With this toolbox, we computed a GLM for each run of each subject, using one predictor for each stimulus type for each subrun. We also included six predictors specifying 3D head motion. Each run's GLM, thus, yielded four t-value activity patterns for each exemplar (one per subrun). Both runs combined yielded eight t-value patterns for each exemplar. We decoded the exemplar (two orientation variants) for each stimulus type with a linear support vector machine (SVM, using the libSVM library - Chang and Lin, 2011) using leave-two-subrun-out cross-validation (Mur et al., 2009). Cross-validation consisted of four folds over which the first, second, third and fourth subrun of both runs were selected as independent test data. We classified stimulus type using all voxels within the quarterfield patch ROIs (gray bars Figure 3a), using only voxels with a radial or vertical preference (red bars Figure 3a) and using only voxels with a tangential or horizontal preference (blue bars Figure 3a). Note that we computed voxel orientation preference based only on the training data during each cross-validation fold. Spurious orientation preferences (resulting from noise) will not replicate in the test data and therefore cannot contribute to significant orientation decoding. Voxel orientation preference was determined for each quarterfield patch by computing the mean difference of t-values between orientations (e.g. radial minus tangential) taking into consideration the patch's receptive-field location. For example, a voxel in a patch representing the right upper visual quarterfield was considered to have a radial preference if t-values were greater for the right tilted than the left-tilted grating (Figure 2). For spirals, a right-upper-field voxel would be considered to have a vertical preference if t-values were greater for the counter-clockwise than for the clockwise spiral.

**Figure 3.**
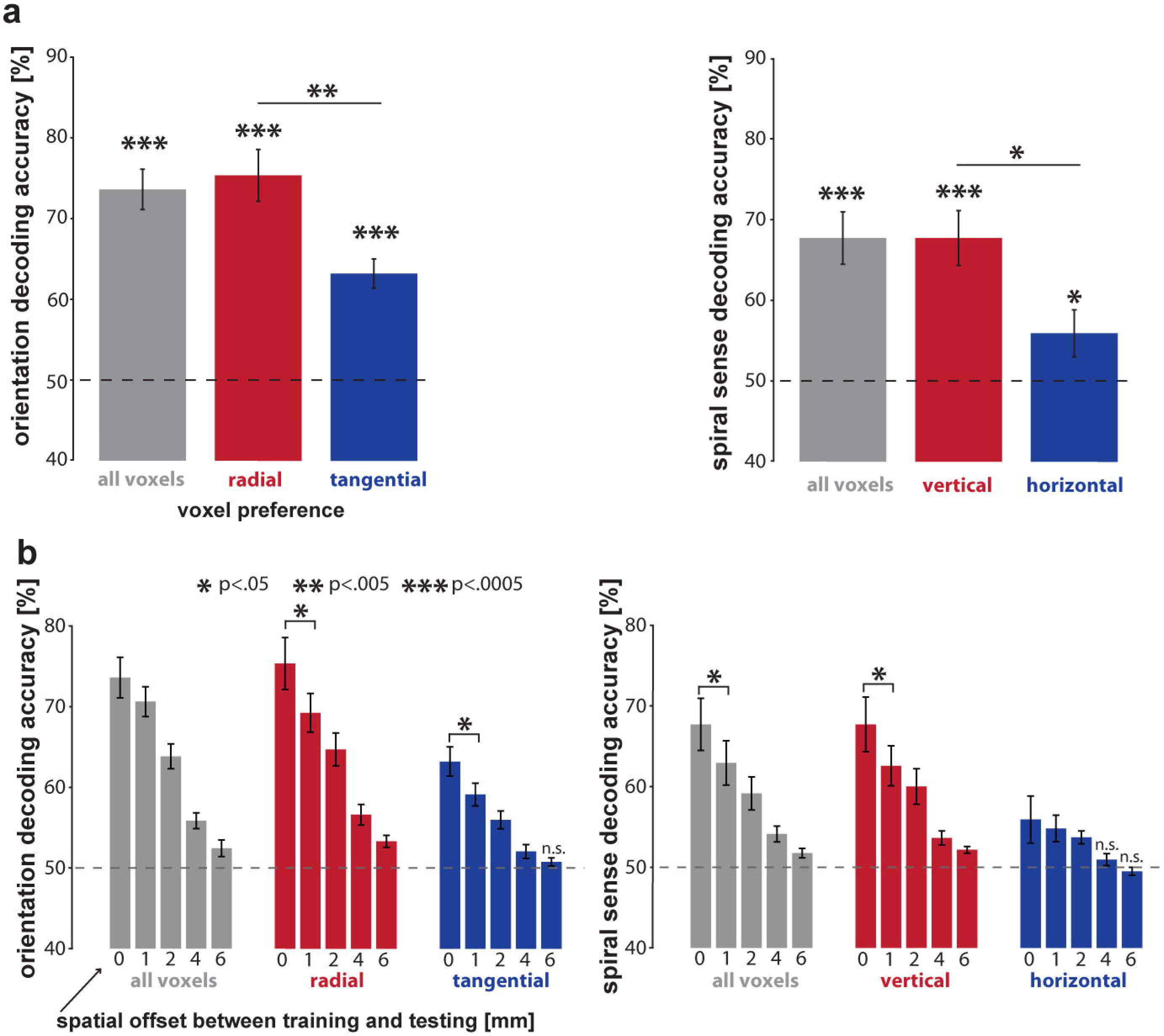
tangential and horizontal orientation preferences are replicable and on their own allow for robust orientation decoding. **(a)** Bar plots summarizing grating orientation (left) and spiral sense (right) decodability when selecting all voxels (gray bars), voxel preferring radial/vertical preference (red bars) and voxel preferring tangential/horizontal preference (blue bars). **(b)** Bar plots summarizing how grating orientation (left) and spiral sense (right) decodability is affected by spatially shifting test patterns by 1, 2, 4 and 6 mm.

*Assessment of the effect of spatial pattern shifts on orientation decodability*. Testing data was spatially shifted by 0.5, 1, 2 or 3 voxels - corresponding to 1, 2, 4 and 6 mm - using shifted ROI coordinates for each patch when computing test patterns. The shift of 0.5 voxel (1 mm) was realized by spatial interpolation (average of two adjacent voxels). Data was shifted in all six directions (ventral, superior, left, right, anterior, and posterior). During this analysis classification performance was computed as the average SVM decoding accuracy across all shift-directions within each participant.

## Results

### Voxels with reliable opposite orientation preferences intermingle within small V1 patches

In order to find out if V1 contains voxels with opposite orientation preferences, we considered the responses to left- and right-tilted gratings in V1 patches at the center of the visual quarterfields (representing polar angles 45°, 135°, 225° and 315° clockwise from vertical; Figure 2a). For a given patch of voxels, each grating thus had either a radial or a tangential orientation. Contrast t maps between the two gratings are shown in Figure 2a for two representative participants. These unthresholded statistical maps suggest that voxels with a radial preference and voxels with a tangential preference intermingle within these patches. However, opposite apparent selectivities might result from the noise in the data. We therefore used decoding analyses of each of the opposite-selectivity voxel sets to assess the reliability of these preferences separately.

First we assessed stimulus decodability using all voxels within a patch together, regardless of the direction of their preference. Consistent with previous studies, decoding analyses using linear support vector machines (Figure 3a) revealed that the two gratings are robustly decodable (74% accuracy, p < 0.0005). The two spirals, similarly, were robustly decodable (68% accuracy, p < 0.0005; Figure 3a). For the gratings, either radial- or tangential-preferring voxels, or both sets might contribute to orientation decodability. For the spirals, similarly, either vertical- or horizontal-preferring voxels, or both sets might contribute. Note that vertical preferences cannot contribute to grating decoding, because the two gratings were balanced about the vertical orientation. Similarly, radial preferences cannot contribute to spiral decoding, because the two spirals were balanced about the radial orientation.

*Tangential- and radial-preferring voxels are intermingled and each set, by itself, supports orientation decoding*. Single voxel responses are noisy. Even if orientation information resulted only from a coarse-scale map of radial preference, we would expect some inverted preference estimates (apparent tangential-preferring voxels), due to the noise in the data. In order to assess whether the tangential preferences were real, we tested their reliability in the decoding framework (Figure 3a). Importantly, orientation preference of voxels was determined independently from the test data. Spurious orientation preferences would not replicate in the test data. We found that grating orientation can be robustly decoded based only on voxels with a tangential preference (63% accuracy, p < 0.0005, one-sided Wilcoxon signed-rank test across subjects). Consistent with the previously reported slight bias in favor of radial preferences (Sasaki et al. 2006; Freeman et al. 2011; Alink et al. 2013), decoding was also possible using only radial-preferring voxels (75% accuracy, p < 0.0005) and the accuracy was significantly greater for the radial-preferring voxel set than for the tangential-preferring voxel set (p < 0.006, two-sided Wilcoxon signed-rank test across subjects). These results show that the two opposite-preference sets of voxels, which are intermingled within the patches of V1, each carry significant orientation information.

*Horizontal- and vertical-preferring voxels are intermingled and each set, by itself, supports orientation decoding*. We performed analogous analyses on the response patterns elicited by the spirals (Figure 3a). We found that spiral orientation can be robustly decoded based only on voxels with a horizontal preference (56% accuracy, p < 0.04, one-sided Wilcoxon signed-rank test across subjects). Again, consistent with the previously reported slight bias in favour of vertical preferences (Mannion et al. 2010; Alink et al. 2013; Freeman et al. 2013), decoding was also possible using only vertical-preferring voxels (68% accuracy, p < 0.0005) and the accuracy was significantly greater for the vertical-preferring voxel set than for the horizontal-preferring voxel set (p < 0.02, two-sided Wilcoxon signed-rank test across subjects). For the spirals, as well, results show that the two opposite-preference sets of voxels, which are intermingled within small patches of V1, each carry significant orientation information.

### V1 voxels exhibit subtle radial and vertical preferences

In order to assess radial and vertical preferences on a group level we estimated the response amplitude difference (in % signal change) between radial and tangential for all voxels across all participants. We plotted the histogram of the radial-tangential response difference across V1 voxels (Figure 2b - left side; pooled across quarterfield patches and the 18 participants). The histogram shows that voxels in V1 are slightly more likely to prefer radial orientations over tangential orientations (56.1% vs 43.9%). These proportions were significantly different (p < 0.0005, two-sided across-subject t-test, test statistic: within-subject %-point difference, subject as random effect). The mean response difference between radial and tangential orientations was 0.038 %-signal-change.

We performed analogous analyses for the spiral stimuli (Figure 2b - right side), which drive each patch with either a vertical or a horizontal orientation. We plotted the histogram of the vertical-horizontal response difference across V1 voxels (Figure 3b; pooled across quarterfield patches and the 18 participants). The histogram shows that voxels in V1 were slightly more likely to prefer vertical orientations over horizontal orientations (58.3% vs 41.7%). These proportions were significantly different (p < .0005, same test as above). The mean response difference between radial and tangential orientations was 0.031 %-signal-change.

Both the radial-over-tangential and the vertical-over-horizontal preference effect sizes were small, less than 2% of the average fMRI response in these V1 patches for gratings and spirals, which were 2.03 %-signal-change and 2.17 %-signal-change, respectively. These results are consistent with the previous analysis of this data set in Alink et al. (2013).

### Shifting test patterns by half a voxel or more reduces orientation decodability

In Figure 2a, we saw that opposite orientation preferences between neighboring voxels intermingle. However, voxels with similar orientation preference appear to form spatial clusters. Freeman et al. (2013) recently found that the decodability of grating orientation and spiral sense is not affected by shifting activation patterns by half a voxel (1 mm) between training and testing. This was taken as evidence for fMRI orientation decoding relying mainly on coarse-scale orientation preference maps rather than intermingled orientation preferences. To relate this finding to our data, we have assessed decoding performance after shifting test patterns by 1, 2, 4 and 6 mm. Our results do show strong effects of spatial shifts on decoding performance (Figure 3b). Even the minimal shift of 1 mm significantly reduced decoding performance for four out of the six voxel selections (Figure 3b, p < 0.05). Orientation decodability for all selections was found to approach chance level for 6 mm shifts. These results are consistent with the presence of information across multiple spatial scales, including fine-grained and coarse-scale patterns.

## Discussion

The aim of the current study was to find out whether fine-grained neural activation patterns contribute to fMRI orientation decoding in the context of acquisition with a typical 3T scanner at a spatial resolution of 2 mm isotropic. Alternatively, fMRI orientation decoding might rely solely on global-areal patterns resulting from radial and vertical preferences. Previous studies used spatial-frequency filtering techniques to address this question. However, spatial-frequency filtering can be confounded by a high-spatial-frequency voxel gain field and can suggest the presence of fine-grained pattern information where there is none (Figure 1, left). Here we exploited the fact that the voxel gain field is not expected to invert the selectivity of a voxel. Therefore, reliable opposite orientation selectivities within a small cluster of voxels indicate fine-grained pattern information (Figure 1, right).

We investigated whether there are two separate sets of voxels in a small patch of V1 that have opposite orientation preference. To ascertain that each set has a reliable preference (and is not just inverted by noise), we performed orientation decoding on each set separately. Our main finding is that grating and spiral orientation can be decoded based on voxel populations with either orientation preference. This finding indicates that global-areal patterns evoked by vertical and radial preference in V1 are not the only source of visual-orientation information in fMRI at 3T and supports the idea that fine-grained activation patterns contribute to orientation decoding.

Our results are also consistent with the previously demonstrated enhanced V1 responses to radial (Sasaki et al., 2006; Mannion et al., 2010; Freeman et al. 2011; Alink et al., 2013) and vertical orientation (Mannion et al., 2010; Alink et al. 2013; Freeman et al., 2013) as voxels preferring vertical and radial orientation were slightly more common than those preferring tangential and horizontal orientation. As expected, we also found that decoding performance was greater when selecting voxels preferring radial and vertical orientation for gratings and spirals, respectively. Our results, thus, replicate the presence of global-areal preferences and demonstrate that fine-grained patterns as well contribute to orientation decoding.

Recently, it has been suggested that that orientation decoding might results from differences in contrast along the edges of annular gratings with different orientations (Carlson, 2014). Contrast along the annular edge of a grating varies as a function of the orthogonality between the grating orientation and local edge orientations. As a consequence, vertical annular gratings give rise to higher contrast at the top and bottom edge while horizontal gratings lead to greater contrast at the lateral edges. This edge effect is thought to give rise to global-areal activation differences similar to those resulting from a radial preference. However, the edge effect does not offer a simple account of spiral decoding, where edge-orientation-contrast patterns are expected to be matched between stimuli (Clifford & Mannion, 2015; Carlson 2015). Therefore, edge effects might contribute a global-areal component to grating decoding, but not to spiral decoding. Moreover, they do not account for the local intermingling of opposite selectivities.

Our finding that fMRI orientation decoding is supported by fMRI voxels with opposite orientation preferences does not imply that these orientation preferences have a salt-and-pepper spatial distribution in V1, or that standard-resolution fMRI reflects subvoxel columnar patterns via a hyperacuity mechanism (Op de Beeck et al., 2010; Kriegeskorte et al., 2010; Boynton 2005; Shmuel et al, 2010). Although voxels are likely to constitute complex spatiotemporal filters (Kriegeskorte et al., 2010), it is an open question whether this supports hyperacuity. Our data shows that voxels preferring the same orientation tend to form clusters within V1 (Figure 2a). The fine-grained orientation preferences we report might result from orientation-specific responses of veins on the scale of the fMRI voxels. Veins might exhibit such a preference because their branches happen to non-uniformly sample columns preferring different orientations. Alternatively, it has been suggested that the vasculature might align itself to the functional architecture of the cortex during development with veins specifically draining from columns of a particular orientation preference (Gardner, 2010).

In summary, we demonstrate that voxels with various preferences intermingle within small voxel clusters. In addition to global-areal patterns resulting from radial and vertical preferences and edge effects, thus, fine-grained patterns do contribute to fMRI decoding at conventional resolution.

## Acknowledgment

This work was supported by the UK Medical Research Council and by a European Research Council Starting Grant (261352) and Wellcome Trust Project Grant (WT091540MA) to NK, a Gates Cambridge Scholarship to AW and a British Academy postdoctoral fellowship to AA.

